# An immune world model with multi-agent orchestration for peptide vaccines

**DOI:** 10.64898/2026.10.19.760096

**Authors:** Letian Chen, Haoyang Luan, Gufeng Yu

## Abstract

Peptide-vaccine design requires a connection between vaccine-induced priming and recognition of the antigen naturally displayed by a tumor. We present an immune world model that preserves this connection through separate vaccine and tumor branches, endpoint-specific evidence and multi-agent orchestration. Native-recognition constraints held an altered peptide despite its higher presentation score, and targeted perturbations removed antigen-dependent benefit when required tumor-recognition conditions were absent. Public dendritic-cell data distinguished donor from antigen-source transfer: a context ranker achieved Recall@20 of 0.483 and 0.333 under the respective holdouts. Structure-informed receptor prediction added candidate-ranking evidence while preserving functional constraints and matched scenario trajectories. An assay observation model further concentrated a cross-presentation coefficient around its known value in a synthetic conditioning experiment. Together, these results establish a computational framework for linking molecular predictions, native-target requirements and conditional immune dynamics. The framework provides a reproducible basis for choosing peptide-vaccine candidates and the experiments needed to resolve their biological dependencies.

## Introduction

Peptide vaccines must connect two biological events: priming T cells against an administered antigen and recognizing the target naturally displayed by a tumor. Antigen-specific responses in personalized vaccination studies, together with the systemic organization of cancer immunity, make this connection a central design problem^1–3^. Changing a vaccine peptide can improve a local molecular score while changing the target that responding cells encounter. A useful computational framework must therefore preserve the relationship between vaccine design and native-tumor recognition throughout the immune pathway.

Existing methods provide complementary parts of this calculation. Sequence predictors estimate peptide presentation or receptor binding^4–6^, whereas integrated immune simulators and quantitative systems pharmacology models connect cell populations, tissue compartments and interventions^7–10^. These outputs describe different endpoints. A presentation score ranks candidate ligands; a receptor score describes a molecular interaction; a dynamical model propagates states under specified rates and initial conditions. Connecting them requires an explicit account of which evidence supports each transition.

Long-peptide vaccination makes this requirement experimentally concrete. Flank and linker design can alter dendritic-cell (DC) presentation^11^, and DC immunopeptidomics measures ligands recovered after uptake of administered material^12^. The tumor, however, processes its endogenous antigen source. Receptor recognition introduces a further dependence on peptide, human leukocyte antigen (HLA) and T-cell receptor (TCR) context, with transfer to unseen epitopes remaining a distinct evaluation problem^13^. The question is how to integrate these local predictions into vaccine comparison while keeping native-target constraints and the evidence needed for downstream inference explicit.

Here we develop an immune world model that separates vaccine and tumor branches and connects them through endpoint-specific evidence. Building on state/action separation in immune world modeling^14^, the framework combines a ten-stage evidence interface, a ten-state mechanistic simulator and five research roles executed by one principal agent. We test its native-recognition constraints, evaluate presentation ranking across donor and antigen-source partitions, integrate structure-informed TCR prediction and condition trajectories on synthetic assay readouts. These evaluations establish a reproducible route from molecular evidence to constrained candidate decisions and conditional immune scenarios.

## Results

### Separate vaccine and tumor branches preserve native-target constraints

The model represents vaccine-side priming and native-tumor recognition as distinct requirements (Fig. 1). A candidate retains its native source, native epitope, vaccine sequence and intended vaccine epitope. These identities connect ten biological stages, from antigen selection and DC uptake to T-cell expansion, killing and memory. Each stage returns an endpoint and its evidence status. This representation allows molecular predictions, measured function and scenario assumptions to contribute to the same candidate decision with their meanings intact.

**Figure 1.**
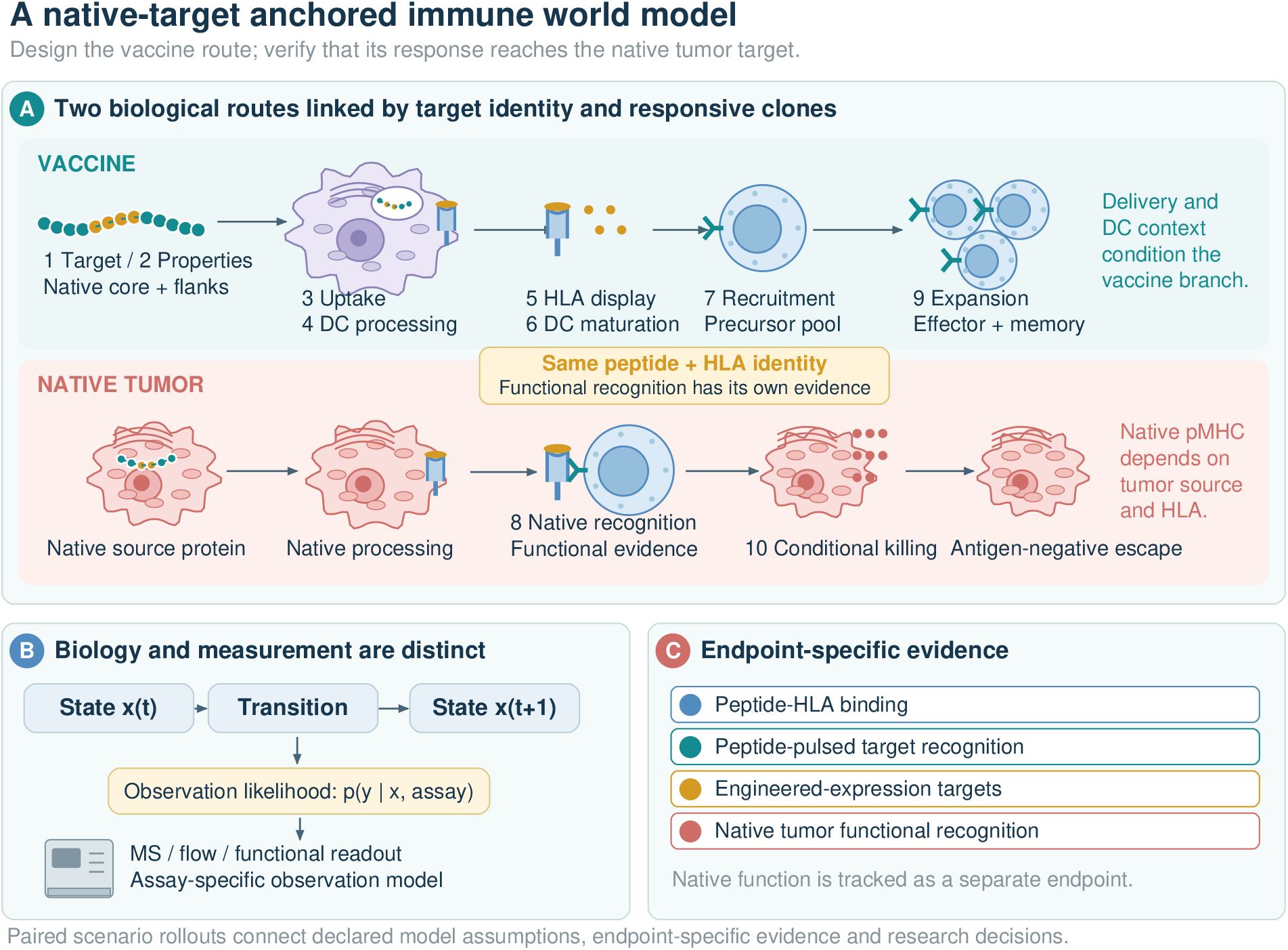
A dual-branch representation of peptide vaccination. (A) Vaccine context determines the administered route through uptake, processing and priming; native antigen and tumor context determine the endogenous target. Numbers identify the ten-stage interface; pMHC denotes a peptide–HLA complex. (B) State transitions and assay observations are represented separately. (C) Binding, peptide-pulsed targets, engineered-expression targets and native tumor recognition retain distinct endpoint labels. The implemented stage map is given in Supplementary Table S5.

The synthetic candidate comparison tested whether a stronger presentation score could bypass native-recognition requirements. An altered core scored 0.627 compared with 0.478 for the native core, but the altered candidate was held because matched vaccine-primed native-functional evidence was absent. Removing recognition, native presentation, tumor HLA or target expression produced exactly zero treated-versus-no-vaccine tumor difference in the tested controls. These perturbations establish that the implemented killing route depends on the specified native-target conditions.

### Presentation ranking resolves distinct donor and antigen-source transfer

To evaluate the molecular evidence entering this framework, we ranked 796 candidate ligands from 12 synthetic long peptides (SLPs) against public DC immunopeptidomics from six donors. Presentation ranking recovered a mean fraction 0.387 of observed ligands in the top 20, compared with 0.341 for affinity ranking (Fig. 2; Supplementary Table S1). The paired gain was 0.046 with a conditional 95% interval of [−0.030, 0.114]. This interval includes zero, and affinity had higher mean recall at a budget of 50. The result therefore supports budget-specific ligand retrieval within this positive-unlabeled candidate pool.

**Figure 2.**
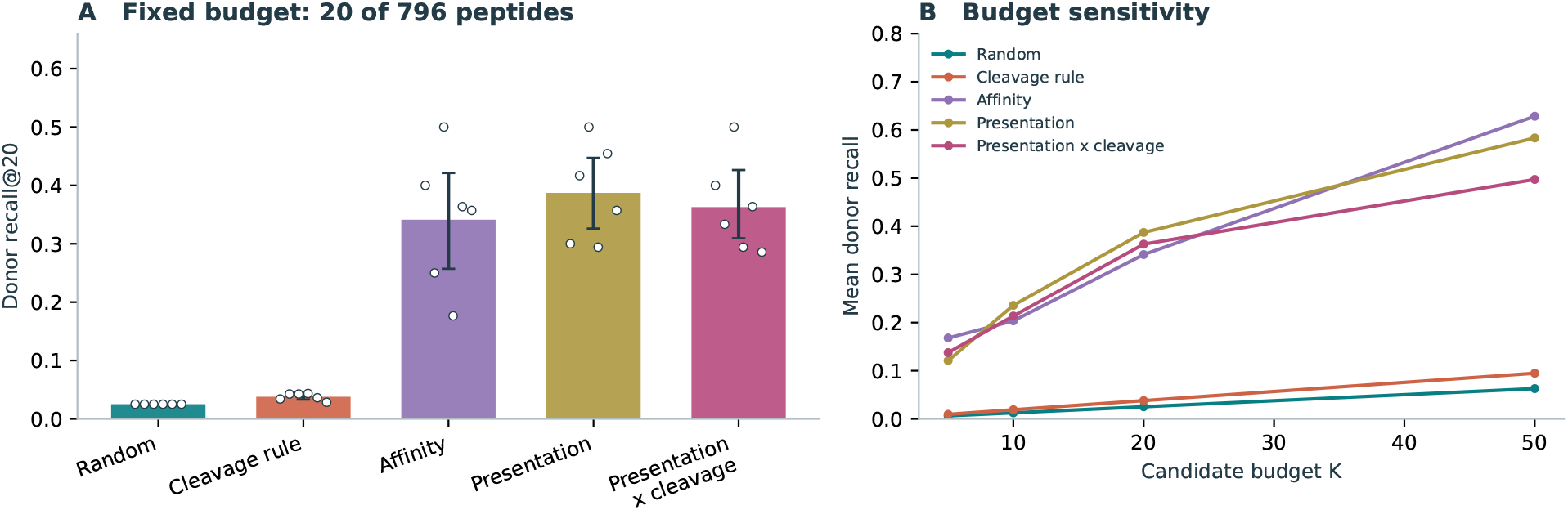
Public DC retrieval. (A) Recall@20 with individual donors shown as white points and conditional bootstrap intervals as error bars. (B) Mean recall across candidate budgets. The endpoint retrieves observed ligands from a positive-unlabeled pool. The presentation-versus-affinity paired interval at *K* = 20 includes zero.

The same evaluation framework also discriminated among proposed scoring updates. Multiplying presentation by the cleavage rule changed mean Recall@20 by −0.024 (95% interval [−0.071, 0.035]). The update did not meet the recorded promotion criteria and was rejected. Retaining this decision connects the adopted scoring rule to the evidence used to choose it.

We then asked whether context features transferred across donors and antigen sources. Across 306 grouped fits, the full context ranker achieved Recall@20 of 0.483 under donor holdout, 0.333 under source-SLP holdout and 0.352 under joint donor–cluster holdout (Fig. 3a; Supplementary Table S2). The fixed pretrained presentation baseline was 0.479 in this donor/adjuvant analysis. HLA-only ranking yielded 0.404 under source holdout, exceeding the full context ranker. Thus donor transfer and antigen-source transfer tested different properties of the fitted model. These evaluations make the relevant transfer axis part of the evidence supplied to candidate design.

**Figure 3.**
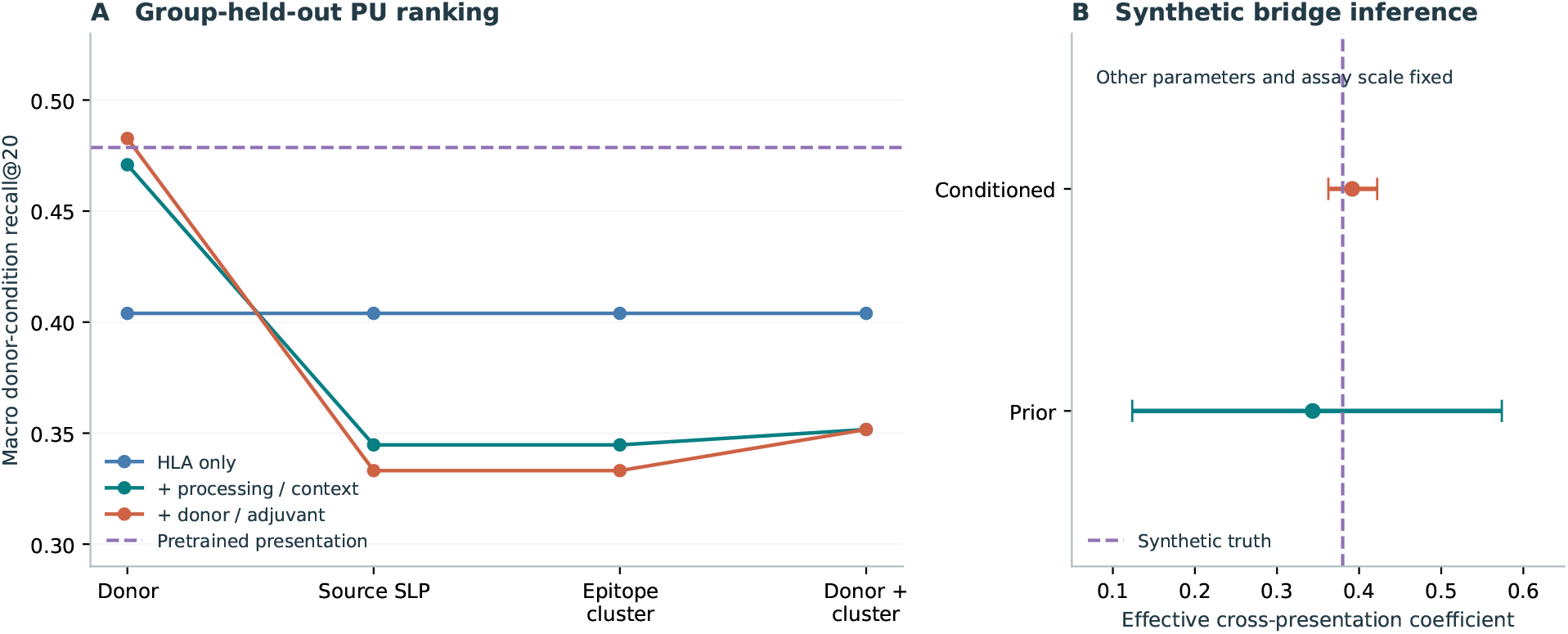
Context transfer and synthetic assay conditioning. (A) Group-held-out positive-unlabeled (PU) ranking. The source and cluster partitions coincide in this dataset. The dashed line is the fixed pretrained baseline. (B) Prior and conditioned 5–95% ranges for one coefficient under synthetic, correctly specified assay data. All other parameters and the assay scale are fixed.

### Receptor prediction adds ranking evidence while retaining functional constraints

StriMap supplied a complementary test of the evidence interface at the receptor endpoint^15^. Three released ensembles returned finite binding-classification scores for 159 distinct queries. In the hypothetical TP53 application, all three candidates received scores for 18 receptor/HLA pairs each. Their native aggregate scores were 0.126, 0.061, 0.055, adding a receptor-ranking objective to candidate comparison.

The matched integration control tested the consequence of adding this endpoint. All 18 candidate/scenario trajectories and their physical couplings were exactly unchanged under shared parameter draws, and altered-epitope gates retained their constraints. Binding predictions therefore enriched the candidate evidence without changing the separately specified native-functional inputs.

Training membership determined the interpretation of the deployment panel. All 19 positive released-example pairs occurred in the training data, so its classification metrics describe deployment diagnostics (Supplementary Table S3). On the selected single-receptor AS8.4 panel, the predefined M2 model had a descriptive score–activation Spearman correlation of 0.565; Base had the higher correlation. Figure 4 presents these readouts alongside TP53 query scores. Measured labels and untested cross-pairs remain separately identified in the accompanying records.

**Figure 4.**
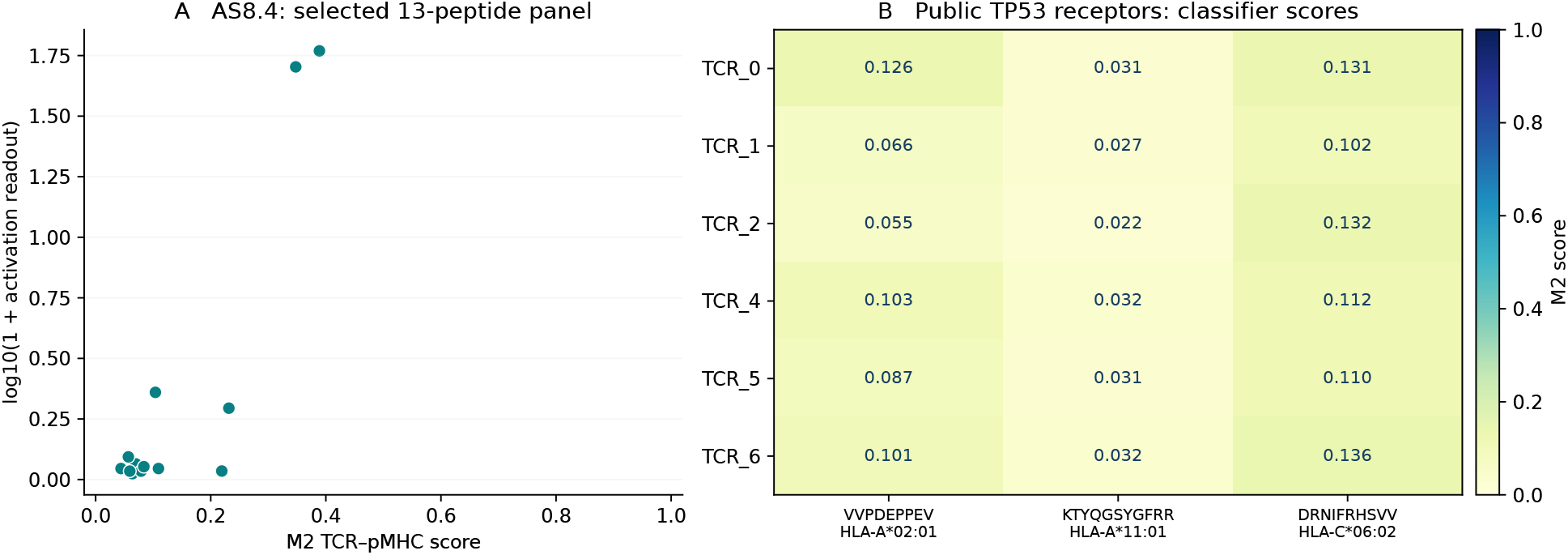
Structure-informed receptor evidence. (A) Released M2 scores versus published activation readouts for 13 selected peptides sharing the AS8.4 receptor; the readout axis uses log_10_(1 + activation). (B) Binding-classification scores for six public TP53 receptors and three peptide/HLA pairs from the hypothetical application matrix. Cross-pairs represent scored queries, with experimental labels retained separately.

### Assay conditioning connects measurements to conditional immune trajectories

The observation model provides a route from endpoint measurements to state inference. In a synthetic experiment, three quantitative readouts concentrated the effective cross-presentation coefficient around its generating value of 0.38. The posterior median was 0.392 with a 5–95% range of [0.363, 0.422] (Fig. 3b). The posterior-to-prior interval-width ratio was 0.132, with an effective sample size of 518.4 among 4,096 particles. All other rates and the observation scale were fixed, making this a controlled test of the conditioning calculation.

Shared parameter draws then connected these mechanistic assumptions to paired intervention scenarios. For the synthetic native long peptide, the median terminal tumor-burden ratio relative to no vaccine was 0.983, with a 5–95% prior range of [0.859, 0.998] (Fig. 5). Low cross-presentation weakened priming, increased suppression reduced the killing coupling and absent recognition removed antigen-dependent benefit. These responses connect the modeled bottlenecks to explicit interventions. Parameter sensitivity and numerical convergence are reported in Supplementary Fig. S2 and Supplementary Note S2.

**Figure 5.**
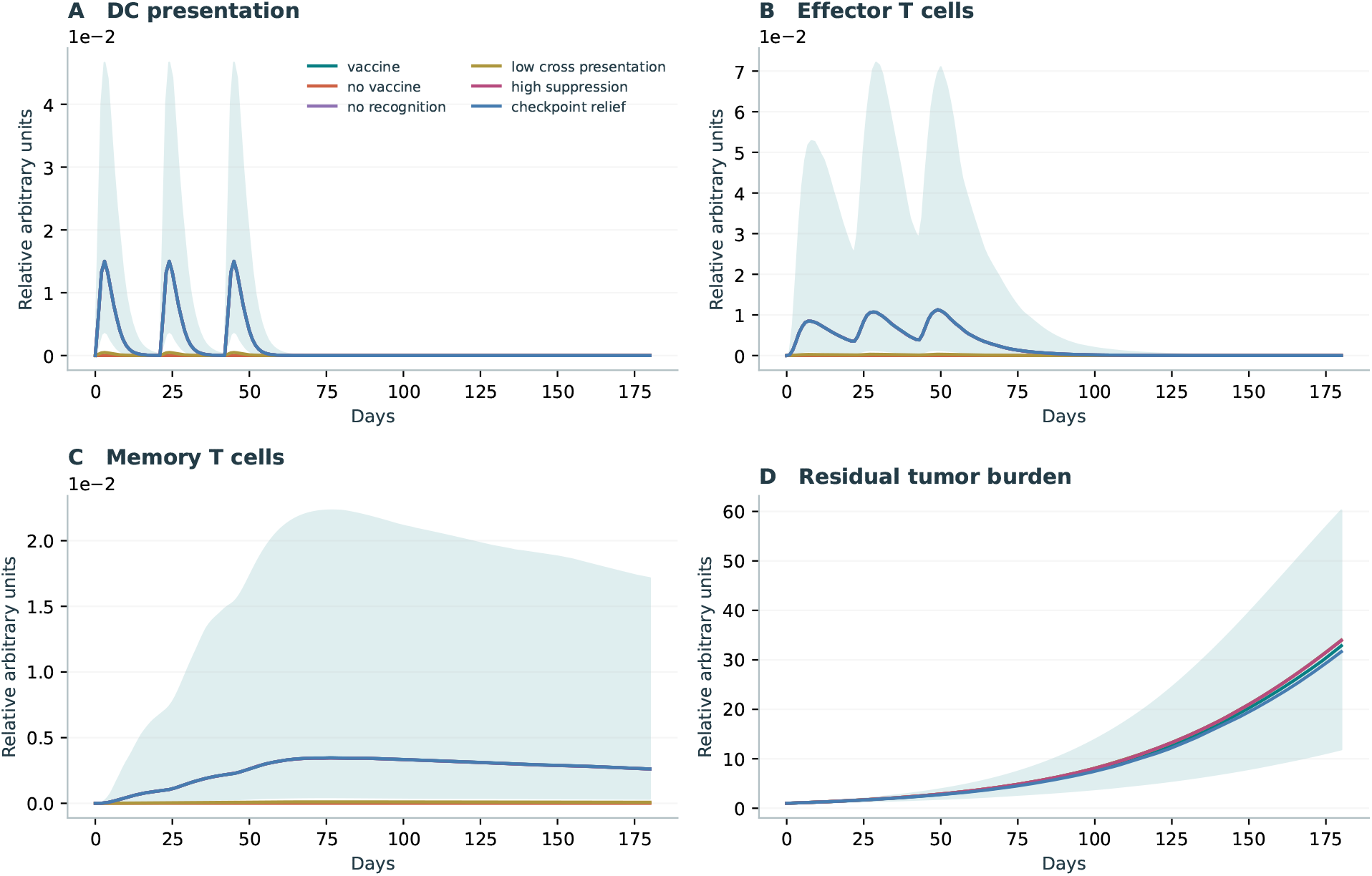
Action-conditioned rollouts for the synthetic native long peptide. Curves show medians over 256 prior draws. The shaded vaccine range is the 5th–95th percentile. (A) DC pMHC proxy, (B) effector T-cell state, (C) memory state and (D) total tumor burden. All magnitudes are relative arbitrary units. Shared draws permit paired scenario comparisons.

## Discussion

This immune world model makes native-target correspondence an explicit constraint on peptide-vaccine research. Its central contribution is a common representation in which local predictions, functional observations and mechanistic assumptions remain identifiable as they enter candidate comparison. The presentation and receptor studies establish the computational interfaces; the perturbation and conditioning experiments establish how those interfaces interact with the declared state model. Together, they support a research workflow that links a proposed vaccine to the evidence required for recognition of its native tumor target.

This organization extends the integration logic of immune simulators and quantitative systems pharmacology^7;9;10^ to an evidence-management problem specific to antigen design. Vaccine-side processing and tumor-side presentation may share an epitope while depending on different biological contexts. Keeping those branches explicit allows an improved local score to be considered alongside the conditions required for tumor recognition. The StriMap integration illustrates the practical value of this separation: receptor information becomes available for ranking while its functional interpretation remains tied to the appropriate observations.

The two molecular evaluations also identify what an informative next experiment must distinguish. Donor and antigen-source partitions probe different transfer questions, while receptor binding and native-tumor assays probe different endpoints. A candidate comparison can therefore point to a specific missing observation: presentation of a new construct, recognition by induced clones, or activity against a native-antigen target. The observation model supplies the computational counterpart by assigning a likelihood to a matched readout and updating parameter weights. Experiment planning can then focus on the biological link responsible for uncertainty in a candidate decision.

The released engine, role prompts and evidence records make this approach reusable. Researchers can replace a molecular predictor, introduce a measured endpoint or compare a mechanistic intervention within the same case representation. The resulting framework supports peptide-vaccine experiments in which candidate identity, evidence scope and downstream assumptions remain traceable from design to interpretation.

### Limitations and future work

The scope of the dynamical inference is set by its calibration. Uptake, cross-presentation, maturation, recognition, tissue effects, memory and escape currently use unfitted priors and relative state scales. The trajectories are conditional simulations; calibrated patient-specific activation, clone expansion, tumor control, toxicity and recurrence require matched longitudinal measurements. The synthetic conditioning study tests one identifiable coefficient with a known observation scale. Jointly unknown abundance and scale can be non-identifiable, and posterior concentration alone does not establish biological validity or predictive coverage.

The molecular evaluations have distinct data boundaries. The DC study uses six donors and a fixed viral SLP pool, with non-detected windows treated as unlabeled. Its bootstrap intervals condition on that pool. Raw spectra were not reprocessed, predictor-training overlap remains unresolved, and the grouped analysis is exploratory on an already inspected dataset. Source and cluster partitions coincide; fold-specific logits are pooled and donor/adjuvant conditions share biological material. Paired alternative flanks of the same core were absent, so these data do not identify the causal effect of a newly designed flank. The cleavage-rule decision applies to the tested rule and endpoint.

StriMap’s example positives all overlap training, leaving no positive class for an independent held-out estimate after exclusion. Exact-identity auditing does not exclude sequence-neighbor, encoder-pretraining or unpublished-data overlap. AS8.4 represents a selected, training-seen receptor evaluated with general released weights. TP53 cross-pairs and mutation decoys lack measured nonbinder controls. Binding-score aggregates therefore remain molecular ranking quantities, separate from repertoire recruitment, functional killing and normal-tissue specificity. The CPU/GPU control tests prediction heads on shared GPU-derived features; timings are workload- and cache-specific.

The ten-stage interface includes explicit unknowns. HLA-II prediction, pMHC stability, population immunogenicity, formulation-dependent uptake and full DC signaling were not validated here. Sequence descriptors and source membership do not establish solubility, in vivo stability or natural tumor processing. The state model aggregates clones and tissue effects, initializes tumor pMHC at a fixed equilibrium and omits spatial structure, evolving proteomics, correlated parameters, process noise, antigen competition and epitope spreading. Measured cell-state transitions, such as those analyzed by CellRank 2^16^, could support later refinements when matched data are available.

Orchestration consists of five sequential roles within one principal agent session. Independent workers, distributed consensus and comparative accuracy or cost benefits were not evaluated. The optional Jev interface exports request contracts and a routing evaluator; hosted calls and a human-labeled routing benchmark were not performed. Candidate design uses constrained enumeration and reranking. Specialist generators, reinforcement learning and new human, animal or wet-laboratory experiments are outside this evaluation.

The next discriminating study is a matched fixed-core construct series measuring DC presentation, induced paired TCRs and responses to native-antigen tumor targets. Time-resolved assays would anchor the presentation scale, while HLA- and antigen-dependency controls and paired normal targets would test the functional link. Receptor- and epitope-held-out panels with measured nonbinders would establish the transfer domain of the binding endpoint. A prospective fixed-budget comparison could then evaluate functional hit rate and assay selection. A separate matched-task study should test orchestration benefits at fixed evidence, biological tools and compute budgets. These experiments would connect the framework’s current evidence contracts to calibrated biological decisions.

## Methods

### Multi-agent orchestration

To coordinate evidence across biological endpoints, the framework separates a decision layer, a biological simulation layer and a data foundation (Supplementary Fig. S1). Five agent roles share a task definition and exchange structured artifacts. The Planner establishes the research question, available inputs, compute budget and comparisons. The Evidence role associates sources and tool versions with biological endpoints and domains. The Design role proposes constrained antigen contexts or interventions. The Critic checks native-target correspondence, evidence coverage, leakage and limiting-case controls. The Experiment role identifies observations that could resolve a candidate decision. A principal agent harness coordinates these responsibilities and calls the deterministic calculation engine.

The released implementation executes the five roles sequentially in one principal agent session, using shared context and role-specific prompts. Here *multi-agent* denotes this separation of research responsibilities. The repository supplies orchestration, stage-decision and criticism prompts, a reusable operating skill and a command-line engine. A case produces validated inputs, stage records, candidate gates, conditional trajectories, reproducibility metadata and an experiment queue. This organization makes each computational action and its evidential basis inspectable.

### Case representation and evidence semantics

A case specifies research context, patient or donor HLA, available TCR information, candidate sequences and an intervention. Each candidate retains four sequence fields: native source, native epitope, vaccine sequence and intended vaccine epitope. Expression, clonality and tumor HLA retention are separate covariates. Schema validation enforces supported residues, exact allele formats, finite values and membership of the native epitope in its supplied source. Synthetic cases and missing observations receive explicit labels.

Each endpoint is labeled as measured, model prediction, heuristic, scenario or unknown. Prediction records retain provider, version, source, endpoint, units and content hash; imported measurements retain assay provenance. Raw-byte hashes connect inputs, code, configurations and outputs. The HTML report and JSON ledger expose the same numerical records, together with evidence coverage and candidate-level decisions.

The ten-stage interface covers target selection; physicochemical properties; delivery and uptake; DC processing; HLA binding and presentation; DC maturation; population immunogenicity; TCR recognition and cross-reactivity; T-cell and tissue dynamics; and killing, memory and regrowth. Supplementary Table S5 maps these stages to implemented calculations and evidence interfaces.

### Molecular predictors and native-target constraints

Target prioritization combines supplied expression and clonality after source-membership checks. Physicochemical calculations include mass, approximate charge at pH 7.4, hydropathy and sequence liabilities. Processing enumerates source-positioned 8–11mer windows. MHCflurry supplies class-I affinity, processing and presentation scores^4;17^, with separate vaccine and native records. A labeled HLA-A*02:01 anchor heuristic supports offline controls. Typed imports additionally accept HLA-II ranks, pMHC half-life and population-immunogenicity endpoints.

Population immunogenicity, clone recruitment and native recognition occupy distinct evidence fields. A qualifying native-functional observation must match candidate, TCR, allele and native epitope, identify vaccine-primed origin, and pass training-record quality checks. Its structured context specifies an unpulsed tumor with naturally occurring antigen expression and HLA- and antigen-dependency controls. Altered epitopes additionally require the matched vaccine epitope. A qualifying negative observation, or an altered core lacking qualifying positive evidence, sets the native-recognition gate to zero. A preserved core uses the declared recognition prior when functional measurements are absent.

Candidate design enumerates source-derived flanks while preserving the native core. Hard constraints precede Pareto ranking within evidence-status groups. The reward vector keeps target identity, DC-presentation proxy, native-functional endpoint, developability, normal-tissue evidence and uncertainty separate. Missing functional endpoints are encoded as null. This constrained enumeration and reranking scheme associates each proposal with both its molecular objectives and its native-target requirements.

### Evidence routing and evaluation contracts

A deterministic classifier maps structured assay fields to binding, peptide-pulsed target, engineered-expression target, native tumor or unknown. Exact identity and numerical thresholds are resolved by code. The repository also supplies a bounded request schema and routing evaluator (Supplementary Note S6); software fixtures exercise the classification contracts.

The feedback interface records endpoint, condition, independent-unit identifier, binary outcome, quality, source and split. Quality-passing training observations update an endpoint-specific Beta(1, 1) model. Held-out observations and duplicate independent units are excluded. A freeze artifact binds raw-byte data and configuration hashes before evaluation.

Each evaluation specifies the endpoint, minimum meaningful gain and a project-specific rationale. Promotion requires a positive lower cluster-bootstrap bound, satisfaction of the gain threshold, zero constraint violations, an independent endpoint and a passed split audit. The cleavage-weighting experiment uses the recorded exploratory threshold of 0.02. Candidate-level confirmation labels are separated from search, and rejected proposals remain in an append-only hash chain.

### Action-conditioned immune dynamics

The state vector is

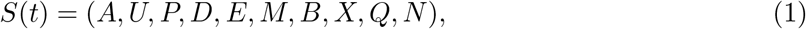

where *A* is extracellular antigen, *U* is DC-internalized antigen, *P* is DC target pMHC, *D* is mature DC fraction, *E* and *M* are effector and memory states, *B* and *X* are antigen-positive and escape-tumor states, *Q* is tumor-native pMHC per normalized antigen-positive cell, and *N* is the available responsive precursor pool. Time is in days; states other than *D* use relative arbitrary units. Interventions specify vaccination times, relative exposure, adjuvant strength and checkpoint relief.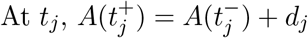. Between events an ordinary differential equation (ODE) defines

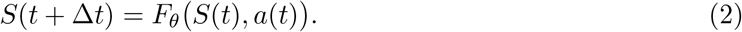

Supplementary Note S4 gives the complete equations and parameter ranges.

Uptake and processing feed DC pMHC through an effective cross-presentation yield. The scenario specification includes a direct-loading path for short peptides and a local rule that halves processing when proline immediately follows the core. Recruitment consumes *N* and depends on DC presentation, maturation, vaccine-side recognition and assumed CD4 help. Native recognition supplies a separate tumor-killing coupling. Responsive clones, trafficking, spatial exclusion and suppression are represented through aggregate states or coefficients.

Tumor populations grow logistically, and antigen-positive cells enter an escape compartment. Antigen-dependent killing requires native presentation, recognition, retained tumor HLA and infiltration; zero supplied expression disables that route. Escape cells have zero killing under the single-target mechanism. Each candidate is simulated independently. Default parameters are independent draws from declared uniform scenario priors, and molecular ranking scores are maintained separately from physical-rate parameters.

Tumor pMHC relaxes toward a native-source/HLA-dependent scenario level and starts at that equilibrium in the demonstration. Flank edits preserve this native level. Antigen, mature DC, effector and memory states start at zero; *N* starts at its sampled precursor pool, and initial burden is partitioned into *B* and *X*. The probabilistic transition is the pushforward of shared parameter draws through deterministic ODEs.

### Observation likelihood and particle conditioning

Biological transition and assay observation are represented separately:

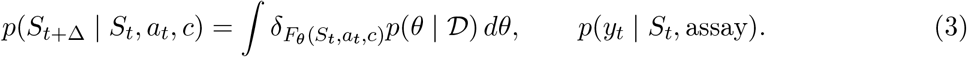

Here *c* is fixed case context, D is compatible conditioning data, and *δ* is a point mass at the deterministic transition. Binary detection uses *p*_det_(*z*) = *f* + (1 − *f*)(1 − *e*^−*ηz*^), with nonnegative latent readout *z*, detection efficiency *η* and false-positive floor *f*. Positive relative readouts use log *y* ~ N (log(*sz*), *σ*^2^) with explicit scale *s*. Both families are configurable assay likelihoods.

Conditioning requires matched candidate, context, endpoint, scale provenance, time, independent unit, split and quality. Static-parameter importance weights satisfy 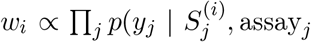 under conditionally independent readout errors. Outputs include weighted trajectories, parameter quantiles and effective sample size 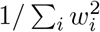 for normalized weights. Low particle support is flagged. Unlabeled MS windows are excluded from conditioning measurements.

The synthetic conditioning experiment estimates the effective cross-presentation coefficient with all other rates and assay scale fixed. Three readouts are generated at days 1, 3 and 7 with log-noise 0.08, generating coefficient 0.38 and 4,096 prior particles. A scale-invariance control checks the equality of likelihoods after compensating changes in latent abundance and assay scale.

Fourth-order Runge–Kutta integration uses event-aligned steps of at most 0.25 day and preserves fractional vaccination times. Non-finite values and negative states beyond roundoff fail execution. Verification repeats selected runs at 0.125 day. Counterfactual scenarios share parameter draws; treated-to-control terminal burden ratios are calculated per draw. Operational regrowth is defined as a decrease of at least 20% below initial burden followed by doubling from the running minimum within the horizon.

### Public DC retrieval experiment

The benchmark uses the official supplementary spreadsheet and PDF of Kessler et al., associated with PXD051490^12^. The source study contains 12 HBV-derived synthetic long peptides (SLPs) and six healthy donors. SLP sequences and HLA assignments were transcribed and visually checked against the supplementary material; peptide lengths were verified. The fixed pool has 796 distinct 8–11mer windows. Seventeen distinct donor alleles yield 13,532 allele–peptide queries.

Length filtering and exact source matching produce 113 donor–adjuvant–peptide records and 25 unique observed sequences, a subset of the source study’s 33 identifications. Isoleucine/leucine ambiguity flags are retained. The primary analysis pools adjuvants within each donor and treats non-detected windows as unlabeled. Each donor’s ranking uses the best score across that donor’s six HLA alleles. Comparators are random selection, a C-terminal cleavage rule, minimum predicted affinity, maximum MHCflurry presentation score, and presentation multiplied by the cleavage rule.

The protocol specifies donor-macro Recall@20 as primary and also reports *K* = 5, 10, 50. Boundary ties are averaged over uniform tie breaking. For observed set *O*_*d*_ and expected inclusion weight 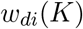,

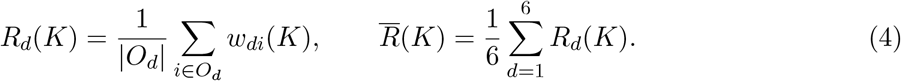

Ten thousand paired donor resamples give conditional bootstrap intervals with the shared SLP pool fixed. The analysis protocol was frozen before inference in this retrospective workflow.

### Grouped context-transfer analysis

The grouped re-analysis retains two adjuvant conditions, producing 9,552 donor–condition–window rows, including 113 detected rows. Three nested feature sets contain HLA affinity alone; affinity plus processing score, fragment length, N/C context lengths and a proline indicator; and these features plus donor/adjuvant indicators. Training-fold standardization precedes a class-balanced logistic ranking surrogate with fixed *L*_2_ penalty 4. Positive-unlabeled (PU) logits rank detected against unlabeled rows.

Leave-one-donor, leave-one-SLP, leave-one-epitope-cluster and joint donor–cluster partitions refit preprocessing and coefficients. Clusters group sources sharing an 8mer with at most one mismatch. The 12 sources form separate clusters, so source and cluster partitions coincide. Joint holdout excludes training rows sharing either held donor or cluster and evaluates their intersection. Across feature sets and partitions this yields 306 fits. Source-containing splits enforce exact peptide disjointness; donor holdout permits shared peptides. Out-of-fold logits are pooled within each donor/adjuvant’s 796 windows, and recall is averaged across 12 conditions. The fixed pretrained baselines retain the same predictions across partitions. Statistical scope and prospective extensions are consolidated in the Limitations and future work section.

### Structure-informed TCR–pMHC integration

Stage 8 integrates the released StriMap TCR–peptide–HLA classifier^15^. Physicochemical features, ESM2 representations and ESMFold monomer geometry enter gated fusion, equivariant graph layers and bilinear attention. The output is a binding-classification score. Inputs require paired CDR3 sequences with explicit V/J alleles or supplied variable domains. Each query is keyed by receptor identity, peptide and exact HLA allele, with HLA-I peptide lengths of 8–11.

Execution uses GitHub revision e14404f and released Zenodo weights. All five pHLA and ten TCR checkpoints per model load strictly. M2, trained by the original authors with peptide-mutation augmentation, was selected before inference as the primary application model; Base and M1 provide fixed comparators. The reported score averages ten classifier outputs using the released parameters.

Three controls preserve input semantics. Structural feature validation aborts on missing or non-finite values. Exact alpha/beta/peptide/HLA length buckets preserve the geometry of real residues during graph inference. Each unique pHLA score and pretrained feature is computed once and reused across receptors, making identical pHLA inputs numerically identical across TCR batches. Head inference disables TF32, and manifests hash encoders, checkpoints, source files, identities and cached features.

The retrospective panel includes 91 released example rows, 13 AS8.4 activation rows, seven published TP53 pairs, and a hypothetical six-receptor by three-peptide by three-HLA application matrix. Deduplication yields 159 queries. An identity audit compares exact CDR3 pairs and peptide/HLA against 22,766 released positive training rows. AS8.4 correlations describe the selected single-receptor panel with general released weights. TP53 examples retain source annotations, with CDR3s derived from motifs in supplied variable domains. One conflicting A*29 annotation is retained as a source-specific query and excluded from the application case and ground-truth metrics.

The same receptor/HLA panel is evaluated against native and vaccine peptides. Stage 8 returns scores, differences, coverage and provenance. Complete coverage enables the mean of the highest up to 30 native scores as an additional Pareto objective. Physical-rate parameters and native-functional gates retain their separate inputs. The application control compares before/after integration using identical priors and random draws.

Numerical controls select the first eight, last twelve and all AS8.4 input rows by identity, giving 33 distinct queries. Batch-size-one GPU and batch-size-16 CPU heads are compared with the full-panel batch-size-16 GPU reference using the same cached GPU-derived features and fixed absolute/relative tolerances 10^−5^*/*10^−4^.

### Computational implementation and verification

MHCflurry 2.2.1 ran with PyTorch 2.8.0+cu128 on an NVIDIA RTX 5090 D. Execution records hash inputs, model files and outputs and record actual CUDA allocation. An agreement control uses 188 demonstration allele–peptide pairs, relative tolerance 10^−4^ and absolute tolerance 10^−6^. Disabling TF32 for matrix multiplication and cuDNN resolved the initial precision discrepancy with the original models and tolerances; both original and corrected records are retained.

Software tests cover schema validation, deterministic replay, ten-stage coverage, no-dose behavior, recognition-null and native-presentation-null cases, HLA loss, absent expression, nonnegative states, fractional events, convergence, altered-epitope gates, feedback partitioning, duplicate units, split leakage, frozen-input integrity, append-only memory and tie-aware retrieval. The synthetic demonstration comprises three candidates, six scenarios per candidate, 256 shared prior draws and a 180-day horizon.

## Data and code availability

The ancillary reproducibility archive, `immune_world_model_code.zip`, contains the calculation engine, role prompts, operating skill, synthetic cases, tests, processed public-data subsets, prediction tables, benchmark results and execution metadata. Its REPRODUCE.md maps the reported results to commands and saved artifacts; a SHA-256 manifest identifies the distributed files. The manuscript source contains editable L^A^TEX, a compiled bibliography and vector figures. Original DC data are available from Kessler et al. and PXD051490^12^. StriMap weights and training data are available at https://zenodo.org/records/18002170; its source supplements accompany Cao et al.^15^. The original software is distributed under the MIT license. Third-party data and models retain their source terms, and model weights are obtained separately. The analysis uses public research data and explicitly synthetic patient contexts.

## Supplementary information

### S1 Research orchestration

Five research roles execute sequentially in one principal agent session. The architecture separates the role workflow, biological state model, assay interface and evaluation contracts. This supplement records the implementation detail supporting the main-text candidate decisions.

**Figure S1.**
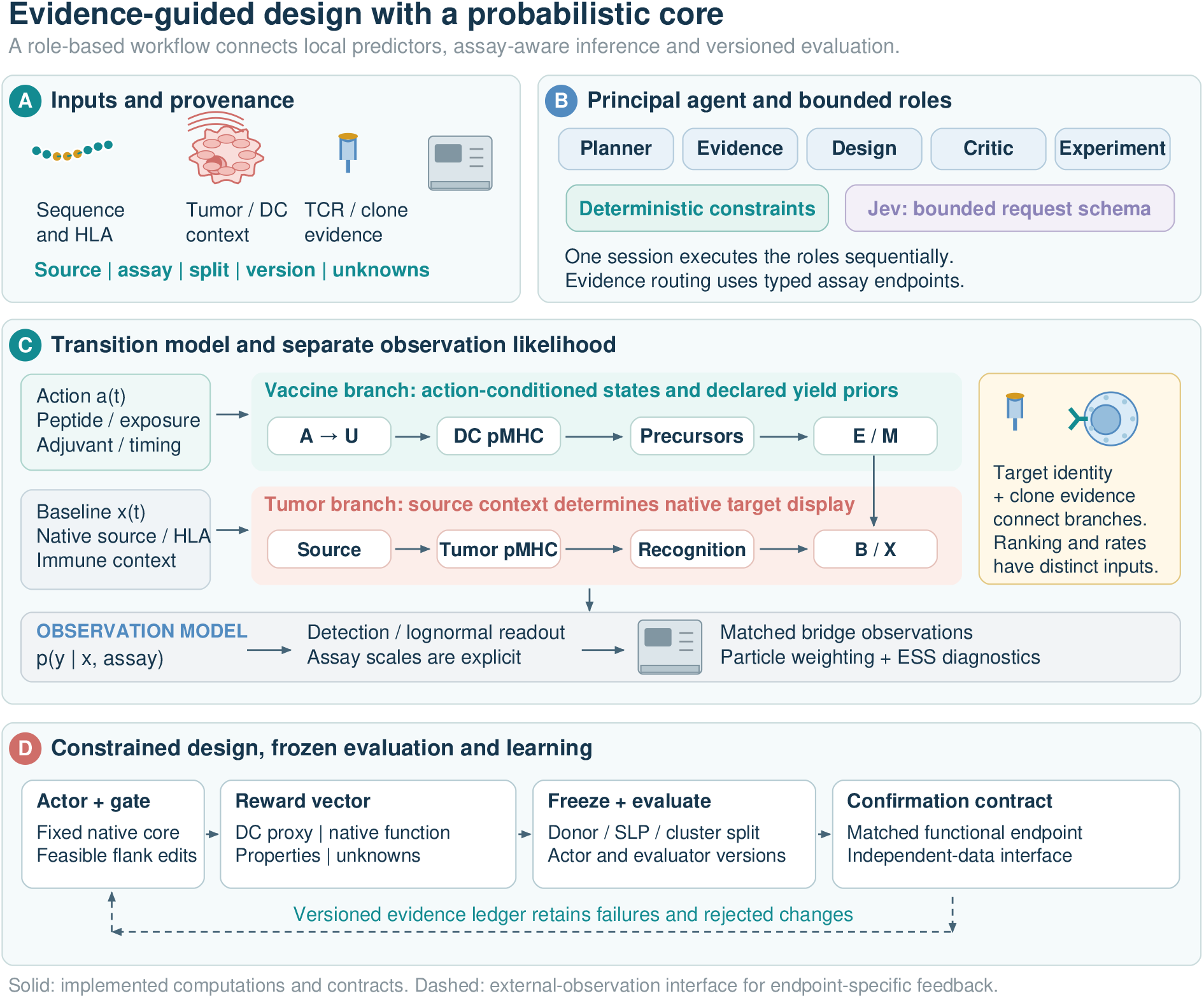
Architecture of the immune world model. (A) Inputs preserve provenance, assay context and data split. (B) One principal agent executes five research roles; code checks exact sequence/HLA constraints, and the optional Jev request schema represents a bounded evidence-routing question. (C) Separate vaccine and tumor branches propagate biological states, while the observation model conditions on matched assays. (D) Candidate constraints, endpoint-specific rewards, frozen evaluation and a confirmation contract organize the design loop. The dashed path denotes the external-observation interface.

### S2 Retrieval and deployment diagnostics

Supplementary Tables S1–S3 retain the complete numerical comparisons summarized in the Results. The six-donor adjuvant-pooled endpoint and the 12-condition grouped endpoint have different aggregation units. StriMap example metrics are deployment diagnostics on training-overlapping positives; AS8.4 correlations are descriptive within a selected single-receptor panel.

All 66 software tests passed. The saved demonstration contains 30 candidate-stage records and 18 trajectories. All terminal states were finite and nonnegative. Halving the integration step gave maximum terminal absolute and relative differences of 0.00258 and 0.00033, respectively, using a 10^−8^ denominator floor. These checks establish consistency with the implemented equations.

**Table S1.** Retrieval at a fixed budget of 20 among 796 candidates. Intervals resample six donors conditional on the shared SLP pool. Random selection is the analytic expectation, not a sampled random baseline.

| Method | Mean donor Recall@20 | Conditional 95% interval |
| --- | --- | --- |
| Random expectation | 0.025 | [0.025, 0.025] |
| Cleavage rule | 0.038 | [0.033, 0.042] |
| MHCflurry affinity | 0.341 | [0.257, 0.421] |
| MHCflurry presentation | 0.387 | [0.326, 0.447] |
| Presentation × cleavage | 0.363 | [0.309, 0.426] |

**Table S2.** Exploratory out-of-fold Recall@20, macro-averaged over 12 donor/adjuvant conditions. Rows denote the entities excluded from fitting. The fixed pretrained presentation baseline is 0.479 in every row. Source and cluster partitions coincide.

| Held-out entities | HLA only | + Processing/context | + Donor/adjuvant |
| --- | --- | --- | --- |
| Donor | 0.404 | 0.471 | 0.483 |
| Source SLP / cluster | 0.404 | 0.345 | 0.333 |
| Donor and cluster | 0.404 | 0.352 | 0.352 |

**Table S3.**
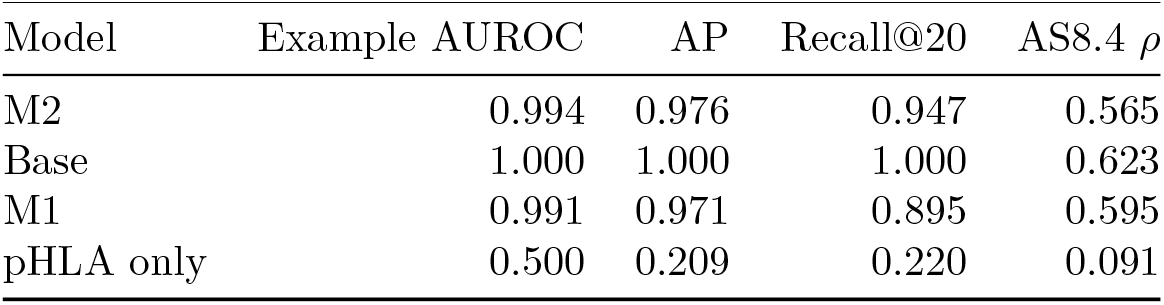
StriMap deployment diagnostics on a training-overlapping example and descriptive AS8.4 correlations. AP is average precision; *ρ* is Spearman correlation for a selected single-receptor panel.

MHCflurry prediction and CSV output for 13,532 queries took 2.08 seconds after model loading, with 139.2 MiB peak PyTorch CUDA allocation. Installation, download, loading and hashing are excluded. In the 188-pair CPU/GPU check, maximum differences were 0.00411 nM for affinity, 1.5 × 10^−7^ for processing and 9.59 × 10^−8^ for presentation. Disabling TF32 resolved the initial discrepancy under unchanged tolerances.

The 159 StriMap queries covered 99 distinct alpha domains, 99 beta domains, 18 peptides and five HLA sequences. Initial M2 execution with a partly populated feature cache took 251.3 seconds and peaked at 29.3 GiB allocation; the final cached execution took 59.6 seconds. The pHLA-only comparator was constant across the released example’s single peptide/HLA and yielded AUROC 0.5. Reusing unique pHLA computations removed an initial batch-dependent difference of 2.4 × 10^−8^ that had broken ties. Final batch-size and CPU/GPU head differences were 1.06 × 10^−7^ and 5.94 × 10^−7^. Original and corrected records are retained.

The complete TP53 matrix, seven source-specific pairs and annotation flags accompany the code. A conflicting A*29 annotation remains a source-specific query and is excluded from application-case and ground-truth metrics. The application scores cover three candidates and 18 receptor/HLA pairs per candidate.

### S3 Prior sensitivity

Descriptive Spearman correlations identify the strongest associations between sampled parameters and terminal tumor burden within the declared independent priors. These sensitivities summarize the assumed model and its parameter ranges. A separate scale-invariance test confirms that compensating changes in latent abundance and assay scale preserve the observation likelihood.

**Figure S2.**
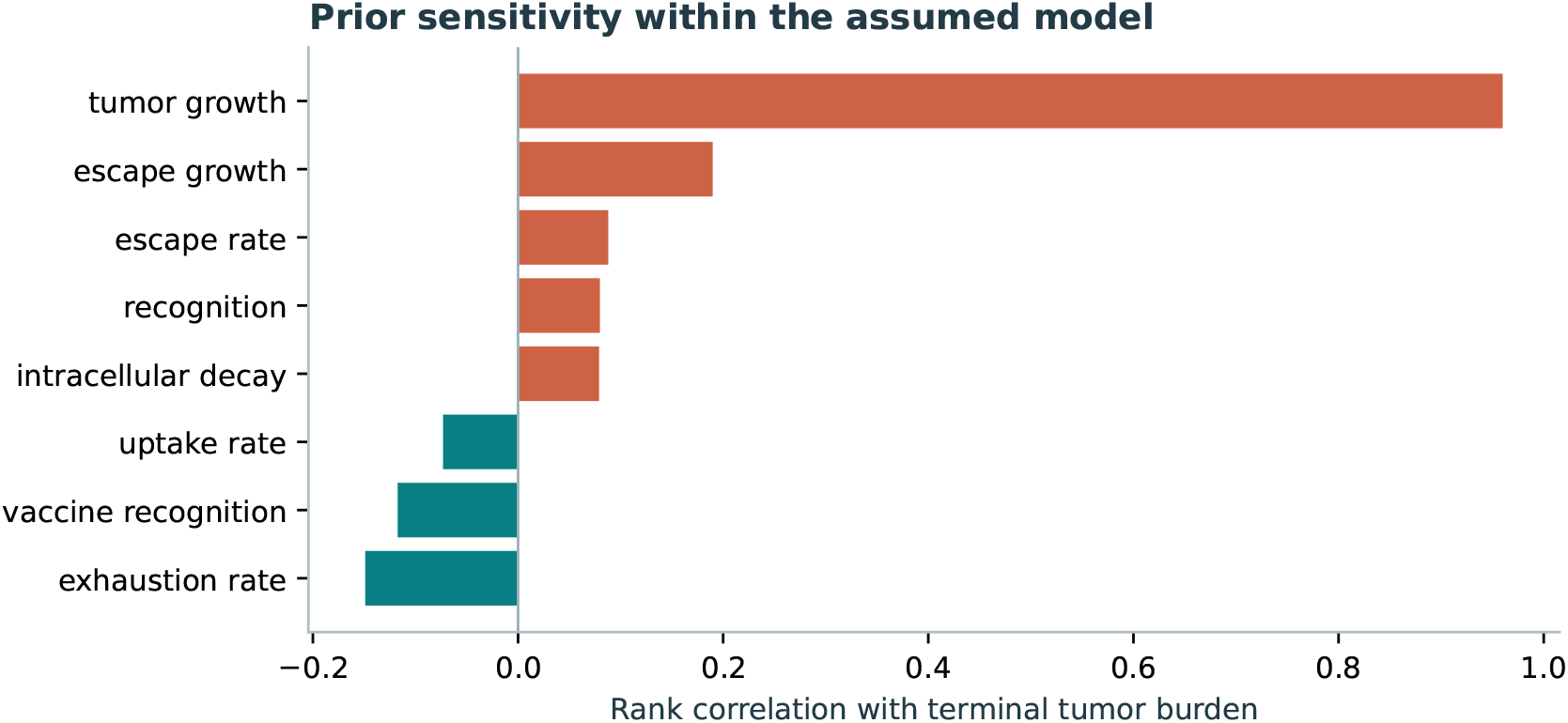
Prior sensitivity within the assumed model. The eight largest absolute rank correlations with terminal tumor burden are shown for the synthetic native long-peptide case. Their interpretation is conditional on the independent prior ranges and equations.

### S4 Transition equations and parameterization

Let *T* = *B* + *X, h*(*P*) = *P/*(0.2 + *P*) and *g*(*T*) = *T/*(1 + *T*). With time in days, the implementation uses

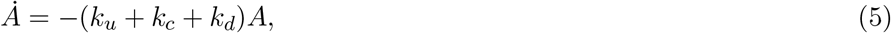

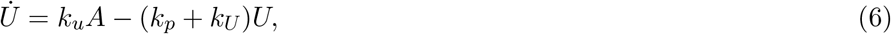

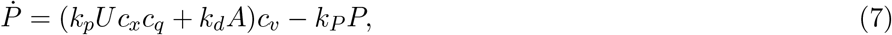

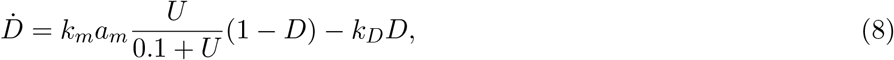

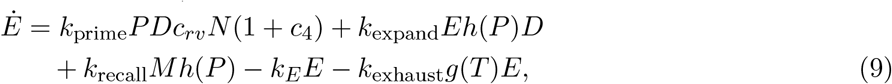

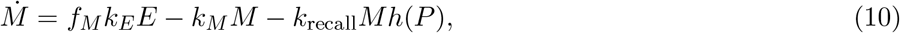

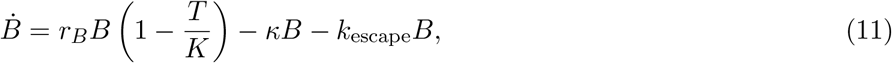

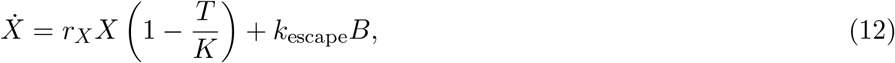

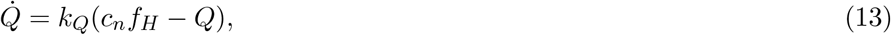

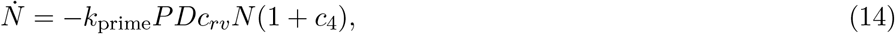

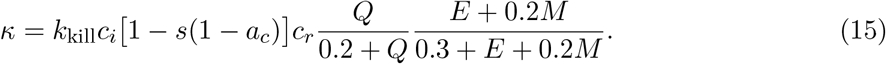

Here *a*_*m*_ is adjuvant strength, *a*_*c*_ is checkpoint relief and *f*_*H*_ is retained tumor HLA, set to zero when supplied expression is zero. The context factor *c*_*q*_ is 0.5 for proline immediately after the vaccine core and 1 otherwise. The direct-loading rate *k*_*d*_ is zero for vaccine sequences longer than 11 residues. Default ranges in Supplementary Table S4 are independently sampled uniform priors. The recognition coefficients *c*_*rv*_ (vaccine-side recruitment) and *c*_*r*_ (native function) are separate. Local predictor scores never multiply *c*_*v*_ or *c*_*n*_. An accepted negative native observation or an unverified altered core sets the native-recognition gate to zero. The coefficients parameterize conditional scenarios.

For the saved demonstration, total initial burden is 1 and carrying capacity *K* = 100. The antigen-positive fraction partitions *B*(0) and *X*(0). Initially *A* = *U* = *P* = *D* = *E* = *M* = 0, *Q* = *c*_*n*_*f*_*H*_ and *N* = *N*_0_. Thus zero antigen input and zero initial vaccine stocks produce no vaccine-driven presentation or recruitment. The default equilibrium *Q* is constant in time. The saturation constants use arbitrary state scales. Recruitment converts precursor units into effector units, but other unlike state scales involve implicit conversion factors. The distributed parameters.json retains exact ranges.

**Table S4:** Default scenario parameter ranges. Relative coefficients are dimensionless. Rate units use the normalized state scales; priming converts pMHC units to effector units.

| Symbol | Source-code parameter | Uniform range | Unit |
| --- | --- | --- | --- |
| $k_u$ | uptake_rate | [0.25, 0.7] | day<sup>-1</sup> |
| $k_c$ | depot_clearance | [0.15, 0.5] | day<sup>-1</sup> |
| $k_d$ | direct_loading_rate | [0.05, 0.2] | day<sup>-1</sup> |
| $k_p$ | processing_rate | [0.25, 0.65] | day<sup>-1</sup> |
| $k_U$ | intracellular_decay | [0.15, 0.45] | day<sup>-1</sup> |
| $c_x$ | cross_presentation | [0.1, 0.6] | relative |
| $k_P$ | presentation_decay | [0.3, 0.8] | day<sup>-1</sup> |
| $k_m$ | maturation_rate | [1, 2] | day<sup>-1</sup> |
| $k_D$ | dc_decay | [0.1, 0.3] | day<sup>-1</sup> |
| $k_{\text{prime}}$ | priming_rate | [0.5, 1.5] | AU<sub>E</sub>/(AU<sub>P</sub> day) |
| $k_{\text{expand}}$ | expansion_rate | [0.3, 0.8] | day<sup>-1</sup> |
| $k_E$ | effector_decay | [0.03, 0.09] | day<sup>-1</sup> |
| $f_M$ | memory_fraction | [0.05, 0.25] | relative |
| $k_M$ | memory_decay | [0.001, 0.005] | day<sup>-1</sup> |
| $k_{\text{recall}}$ | memory_recall | [0.02, 0.08] | day<sup>-1</sup> |
| $k_{\text{kill}}$ | killing_rate | [0.1, 0.35] | day<sup>-1</sup> |
| $r_B$ | tumor_growth | [0.012, 0.03] | day<sup>-1</sup> |
| $r_X$ | escape_growth | [0.012, 0.03] | day<sup>-1</sup> |
| $k_{\text{escape}}$ | escape_rate | [0.0001, 0.001] | day<sup>-1</sup> |
| $k_{\text{exhaust}}$ | exhaustion_rate | [0.01, 0.06] | day<sup>-1</sup> |
| $s$ | suppression | [0.2, 0.7] | relative |
| $c_i$ | infiltration | [0.3, 0.9] | relative |
| $c_r$ | recognition | [0.1, 0.8] | relative |
| $c_n$ | native_presentation | [0.1, 0.8] | relative |
| $c_v$ | vaccine_presentation | [0.1, 0.8] | relative |
| $c_4$ | cd4_help | [0, 0.5] | relative |
| $c_{rv}$ | vaccine_recognition | [0.1, 0.8] | relative |
| $N_0$ | precursor_pool | [0.2, 1] | relative |
| $k_Q$ | tumor_pmhc_turnover | [0.2, 0.6] | day<sup>-1</sup> |

### S5 Ten-stage capability and evidence map

**Table S5:** Implemented calculations and evidence interfaces across the ten biological stages. Each output retains its endpoint and evidence label.

| Stage | Biological question | Implemented interface and output |
| --- | --- | --- |
| 1 | Target and candidate selection | Source membership, expression/clonality priority and native-core context enumeration. |
| 2 | Physicochemical properties | Mass, approximate charge, hydropathy and sequence-liability descriptors. |
| 3 | Delivery and DC uptake | Antigen-depot, uptake and clearance trajectories under declared priors. |
| 4 | Processing and cross-presentation | 8–11mer windows, MHCflurry processing output and explicit cleavage control. |
| 5 | HLA binding and presentation | MHCflurry class-I scores, separate vaccine/native records and typed endpoint imports. |
| 6 | DC maturation and adjuvant effects | Adjuvant-conditioned mature-DC state within the scenario model. |
| 7 | Population immunogenicity | Independent endpoint import and evidence-status contract. |
| 8 | Specific recognition and cross-reactivity | Released StriMap binding scores, matched functional observations and altered-epitope constraints. |
| 9 | T-cell and tissue dynamics | Effector expansion, attrition, recall and aggregate suppression/infiltration. |
| 10 | Killing, memory and regrowth | Antigen-dependent killing, escape, memory and operational-regrowth trajectories. |

### S6 Optional evidence-routing interface

The software can export a bounded assay-classification question using the Jev request format. Its documented Choice/Score confidence statistics describe answer distributions^18;19^. A separate evaluator accepts externally labeled predictions and reports classification error, multiclass Brier score and risk–coverage at a specified threshold. Software fixtures exercise these interfaces, with their evaluation scope described in the Limitations and future work section.

### S7 Reproducibility and figure provenance

Performance estimates, numerical-agreement errors and main rollout summaries are inserted from saved benchmark, rollout, GPU and verification artifacts through values.tex. Dataset counts and model settings are checked against the corresponding records and source code. The ancillary code archive includes donor-level retrieval values, prior-sensitivity data, computation scripts and prediction tables. Original failures remain alongside the corrected CPU/GPU comparison and final StriMap predictions. In particular, the initial StriMap smoke check identified a missing temporary PDB directory in the upstream local-checkpoint path; creating that directory enabled structure generation without replacing missing features by a sequence-only fallback. The initial pHLA batching artifacts are retained separately from the final benchmark.

Figure 1 and Supplementary Fig. S1 were first designed using an image-generation tool, then recreated as editable SVG elements and exported to vector PDF with corrected terminology, endpoint scope and arrow direction. No generated raster is embedded in the final vector diagrams. Numerical figures derive from saved artifacts. The ancillary materials preserve the editable SVG sources and calculation evidence; the manuscript source includes the vector PDFs required for compilation. Standard pdfL^A^TEX and BibTeX compile the paper without network access or shell escape.

